# Impact of secondary TCR engagement on the heterogeneity of pathogen-specific CD8^+^ T cell response during acute and chronic toxoplasmosis

**DOI:** 10.1101/2022.01.25.477653

**Authors:** Lindsey A Shallberg, Anthony T Phan, David A Christian, Joseph A Perry, Breanne E Haskins, Daniel P Beiting, Tajie H Harris, Anita A Koshy, Christopher A Hunter

## Abstract

Initial TCR engagement of CD8^+^ T cells results in T cell expansion, and these early events influence the generation of diverse effector and memory populations. During infection, some activated T cells re-encounter cognate antigen, but how these events influence local effector responses or formation of memory populations is unclear. To address this issue, OT-I T cells which express the Nur77-GFP reporter of TCR activation were paired with *T. gondii* that express OVA to assess the impact of TCR activation on CD8^+^ T cell responses. During acute infection, TCR stimulation in affected tissues correlated with parasite burden and was associated with markers of effector cells while Nur77-GFP^-^ OT-I showed signs of effector memory potential. However, adoptive transfer of Nur77-GFP negative or positive OT-I from infected mice into naive recipients resulted in formation of similar memory populations. During the chronic stage of infection in the CNS, TCR activation was associated with large scale transcriptional changes and the acquisition of an effector T cell phenotype as well as the generation of a population of CD103^+^ CD69^+^ Trm like cells. However, while inhibition of parasite replication resulted in reduced effector responses it did not alter the Trm population. These data sets highlight the contribution of recent TCR activation on the phenotypic heterogeneity of the CD8^+^ T cell response but suggest that this process has a limited impact on memory populations at acute and chronic stages of infection.

**Author Summary:** CD8^+^ T cells are important to control many acute and chronic infections, however the role that recent T cell receptor stimulation plays in the formation of ongoing T cell responses is unclear. Here, we utilize a genetic reporter of TCR stimulation and high parameter flow cytometry to characterize TCR-driven phenotypes of pathogen specific T cell responses to the parasite *Toxoplasma gondii* during acute and chronic infection in the periphery and central nervous system. This work demonstrates the importance of recent TCR stimulation in driving local effector CD8^+^ T cell responses in peripheral tissues during infection, as well as the plasticity of the formation of memory T cells. Additionally, we utilize static and live imaging to investigate how *Toxoplasma gondii* life cycle impacts the ability to present antigen to CD8^+^ T cells. These studies aid in our understanding of how effector and memory CD8^+^ T cell responses are generated and maintained during an infection.

## Introduction

CD8^+^ T cells play an important role in immunity to intracellular infections and multiple subsets including T effector, T memory, and T resident memory (Trm) cells have been associated with different anatomical locations and distinct functions [1–3]. During infection, dendritic cells in secondary lymphoid organs present microbially derived peptides via MHC Class I, activating T Cell Receptor (TCR) signaling of naive CD8^+^ T cells. This signal 1, along with co-stimulation (signal 2) and cytokines (signal 3) induce a proliferative burst of CD8^+^ T cells that results in 1,000 to 10,000 daughter cells produced from an individual CD8^+^ T cell within a week of stimulation, which gives rise to diverse effector and memory populations [4–10]. After initial priming, CD8^+^ T cells leave the secondary lymphoid organs and migrate to the periphery, where they can mediate TCR-dependent effector functions that include cytolysis of infected cells and secretion of cytokines (IFN-γ and TNF-α) required for resistance to infection. As pathogen burden is controlled there is a contraction phase where the majority of effector CD8^+^ T cells undergo apoptosis, but a heterogeneous population of long-lived memory T cells remain in the circulation and reside in tissues [6, 11, 12]. During many chronic infections, a pool of effector and memory CD8^+^ T cells will persist that contribute to the long-term control of the pathogen [13–16]. Multiple studies have documented heterogeneity in the phenotype, function, longevity, and fate of these pathogen specific CD8^+^ T cells [8, 17, 18], but it is unclear whether re-encounter with cognate antigen at local sites of infection contributes to the observed diversity.

In current models, antigen experienced T cells are less dependent on signal 2 and 3, and stimulation through the TCR is considered the critical signal that induces local effector activities [19, 20]. However, previously activated T cells can respond to signals 2 and 3 to mediate effector functions that include the production of cytokines such as IFN-γ [21–23]. The ability to assess T cell function and activation status at sites of infection and inflammation has depended on changes in the expression of surface proteins such as CD69, CD44 and LFA-1, the use of genetically encoded reporters of cytokine production or the ability to secrete cytokines on re-stimulation, and even intravital imaging of their behavior [24–26]. These approaches have been complimented by protein and transcriptional profiling that have highlighted phenotypic and functional heterogeneity and how functionality and phenotype of CD8^+^ T cells varies with different pathogens [1, 27, 28]. Nevertheless, certain combinations of surface molecules, such as chemokine receptors as well as activating or inhibitory receptors have helped define different T cell subsets. In certain viral and bacterial infections, the use of graded expression of the chemokine receptor CX3CR1 along with KLRG1 has been used to distinguish effector and memory CD8^+^ T cell populations [8, 9]. Likewise, during toxoplasmosis, the co-expression of cell surface receptors KLRG1 and CXCR3 distinguishes CD8^+^ T cell subsets that have memory potential or are terminally differentiated, with continued presence of infection driving the KLRG1^+^CXCR3^-^ terminally differentiated population [18, 29].

As a consequence of T cell activation, changes in the levels of surface markers such as LFA-1, CD44, and CD69 have been useful to identify antigen experienced T cell populations, but it has been a challenge to distinguish those T cell populations that have undergone recent TCR mediated activation [30]. Thus, while TCR signals promote a marked increase in CD69, this transmembrane C type lectin associated with tissue retention is also upregulated by inflammatory signals, its downregulation is varied and following clearance of antigen CD69 is constitutively expressed on Trm populations [31–34]. Genetically encoded reporters of calcium flux have been useful for imaging of TCR signals *in vivo* [24], however these calcium fluxes are rapid and transient and this method does not permit long term tracking or isolation of antigen stimulated cells. Nur77 is an orphan nuclear hormone receptor that is upregulated quickly following stimulation of the TCR [35–38] and transgenic mice that express Nur77-GFP have been used to identify T cells that have experienced recent TCR engagement during priming and the effector phase of T cell responses [35, 36, 39].

*Toxoplasma gondii* is a ubiquitous obligate intracellular parasite that naturally infects many intermediate hosts, including both humans and mice [40]. Resistance to *T. gondii* is dependent on cell mediated immunity and as a natural host for this pathogen, the mouse has provided a tractable model that helped established the role of IFN-γ and CD8^+^ T cells in resistance to intracellular infection [14, 18, 41–43]. The early phase of infection is characterized by high levels of replication of the tachyzoite stage of the parasite and vascular dissemination to multiple tissues [44–46]. This phase of infection generates a robust T cell response that restricts parasite growth, but the ability of *T. gondii* to form latent tissue cysts in neurons allows infection to persist in the CNS [43, 47]. However, in patients with acquired defects in T cell responses, the reactivation of the tissue cysts can lead to the development of toxoplasmic encephalitis (TE) [40, 48]. This is recapitulated in mice where long-term control of *T. gondii* is mediated by CD4^+^ and CD8^+^ T cell production of IFN-γ which activates a variety of anti-microbial mechanisms in hematopoietic and non-hematopoietic cell types [49–51]. Analysis of the CD8^+^ T cell responses during acute and chronic toxoplasmosis has highlighted mechanisms that generate and sustain CD8^+^ T cell responses in the periphery [29, 52–54] as well as the presence of a sub-population of Trm-like cell populations in the CNS [18, 33, 52] that are mirrored in other infections [11, 55].

The ability to modify *T. gondii* to express model antigens such as ovalbumin (OVA), combined with TCR transgenics has provided an important tool to understand T cell responses to this infection [56–60]. This approach allowed the study of early events that lead to parasite specific T cell responses and how these operate in the CNS [18, 61, 62]. Here, OT-I CD8^+^ T cells specific for the OVA peptide SIINFEKL and expressing Nur77-GFP were combined with *T. gondii*-OVA to identify cells that had recently engaged their TCR and asses how this impacted function. Although TCR activity has been linked to memory formation [63], an adoptive transfer model indicated that during the acute phase recent TCR stimulation did not influence the ability to form long lived memory T cells. During the chronic stage of infection, tachyzoite replication in the CNS was associated with TCR activation, large scale transcriptional changes, and the acquisition of an effector T cell phenotype as well as the generation of a local population of Trm like cells. However, while inhibition of parasite replication resulted in reduced effector responses it did not alter the Trm population. These data sets highlight the contribution of recent TCR activation on the phenotypic heterogeneity of the CD8^+^ T cell response but suggest that T cell memory formation and maintenance is independent of recent TCR activation.

## Results

### Nur77-GFP reporter detects recent *T. gondii*-mediated TCR activation

To track TCR activation in a defined population of CD8^+^ T cells, transgenic mice were generated that express the OT-I TCR specific for the H2-Kb restricted SIINFKL peptide and the Nur77-GFP reporter (Nur77-GFP OT-I). As expected, naive Nur77-GFP OT-I stimulated with αCD3 *in vitro* induced expression of GFP within 45 minutes that peaked by 2 hours (Figure 1A). Removal of αCD3 stimulation resulted in downregulation of GFP expression within 24-36 hours with basal levels by 72 hours (Figure 1B). This stimulation was accompanied by expression of CD69 which persisted after T cells were rested and had lost GFP expression (Figure 1C). When rested OT-I were restimulated with αCD3 there was rapid re-expression of Nur77-GFP (Figure 1C). The use of peptides with differing affinity for the OT-I TCR revealed a hierarchy wherein a low affinity peptide (EIINFEKL) did not induce reporter expression, moderate affinity peptides (SIIVFEKL and SIIQFEKL) induced intermediate levels of GFP, and the highest levels were associated with SIINFEKL (Supplementary Figure 1A). To test whether Nur77-GFP OT-I encounter with endogenously processed OVA from an infected cell (as opposed to peptide pulsed cells or αCD3) would activate the reporter, bone marrow derived macrophages were infected *in vitro* with a non-replicating strain of *T. gondii* (CPS or CPS-OVA) and co-cultured with naive Nur77-GFP OT-I. With CPS-infected cells there was no induction of GFP and a modest impact on CD69, but co-culture with CPS-OVA infected cells resulted in high levels of GFP and co-expression of CD69 (Figure 1D). To determine if TCR-independent inflammatory signals during infection influence T cell expression of Nur77-GFP, pre-activated and rested Nur77-GFP OT-I T cells were transferred into mice infected with the PRU strain of *T. gondii* or PRU expressing OVA (*T. gondii*-OVA). Although transfer of these T cells into mice infected with *T. gondii* that did not express OVA resulted in a modest increase of CD69, GFP expression was only observed when OT-I were transferred into *T. gondii*-OVA infected mice (Figure 1E). Thus, in the absence of cognate antigen the inflammatory environment is not sufficient to induce reporter expression. These *in vitro* and *in vivo* data sets validate that the expression of Nur77-GFP provides a sensitive and specific indicator of the recent interaction of OT-I CD8^+^ T cells with *T. gondii*-derived SIINFEKL.

**Fig 1.**
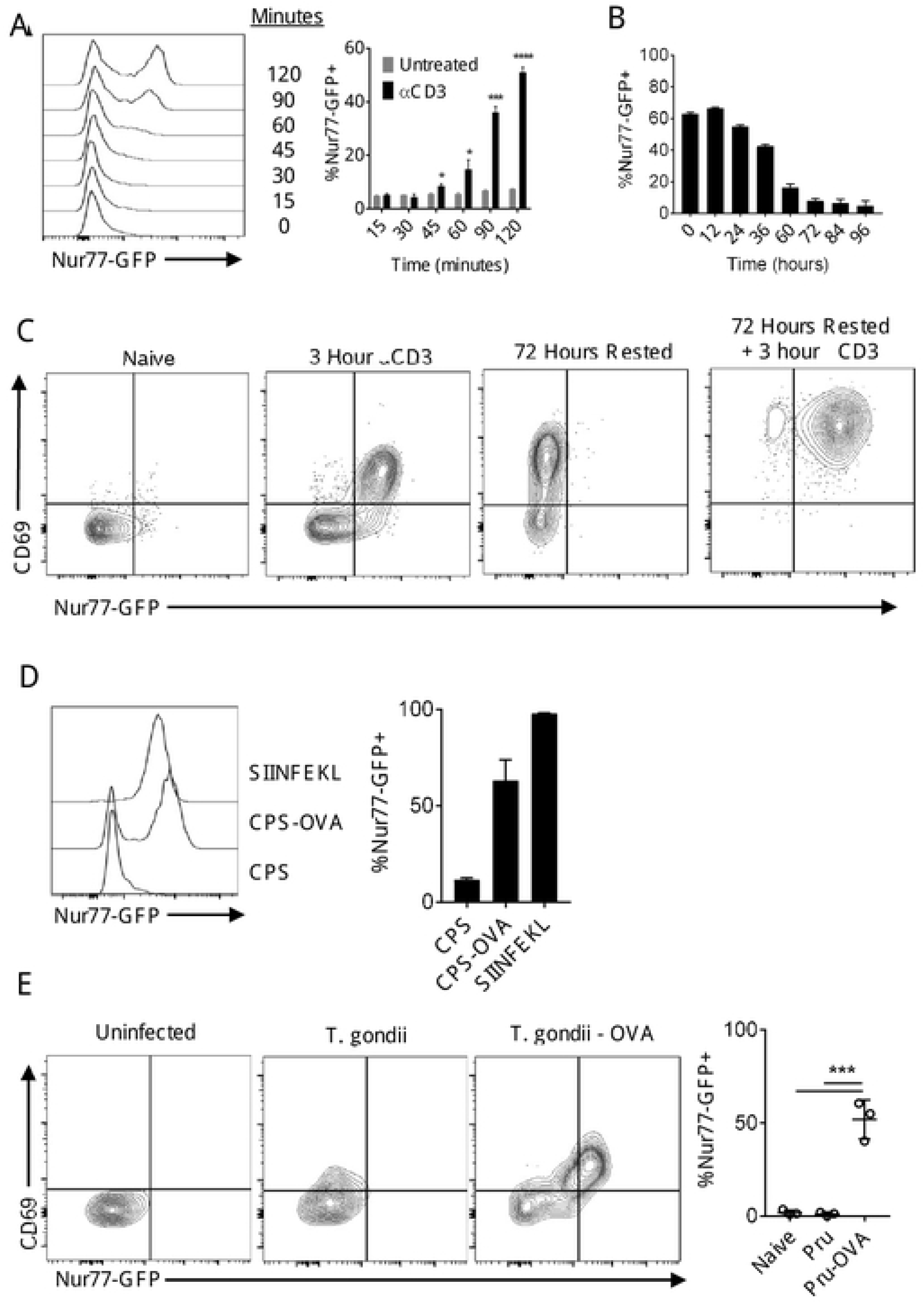
Nur77-GFP OT-I respond to *T. gondii* antigen. (A-B) Naive OT-I stimulated *in vitro* with plate bound αCD3 and αCD28 for up to 48 hours, then removed from stimulation. (C) Rested OT-I restimulated with αCD3. (D) BMDM infected with CPS or CPS-OVA and co-cultured with OT-I. (E) Splenic OT-I following infection with *T. gondii* or *T. gondii*-OVA (N=3 mice per group). P values based on Students T-test with Bonferroni-Dunn correction for multiple comparisons; error bars indicate SEM.

### Infection induced kinetics and phenotype of *T. gondii* specific CD8 T cells

Previous studies have shown infection with *T. gondii*-OVA results in the rapid clonal expansion of OT-I that is dependent on the presence of OVA [28, 56]. To assess the frequency with which these primed T cells re-encounter antigen, 5,000 congenically distinct (CD45.1) naive Nur77-GFP OT-I were transferred into CD45.2 recipients that were then infected with *T. gondii*- OVA that expresses tdTomato. To focus on TCR mediated events after initial priming, mice were analyzed between 10 and 60 days post-infection (dpi). The number of live cells in the spleen that were positive for tdTomato revealed high parasite burden at 10 dpi with a marked decline by days 14 and 20 (Figure 2A). Numbers of OT-I peaked at 14 dpi and the contraction was apparent by 20 dpi (Figure 2B). Mirroring pathogen burden, at 10 dpi GFP^+^ OT-I in the spleen were readily detected with peak numbers at day 14 (Figure 2B). As parasite burden declined, there was a marked reduction in the numbers and frequency of GFP^+^ OT-I (Figure 2B).

**Fig 2.**
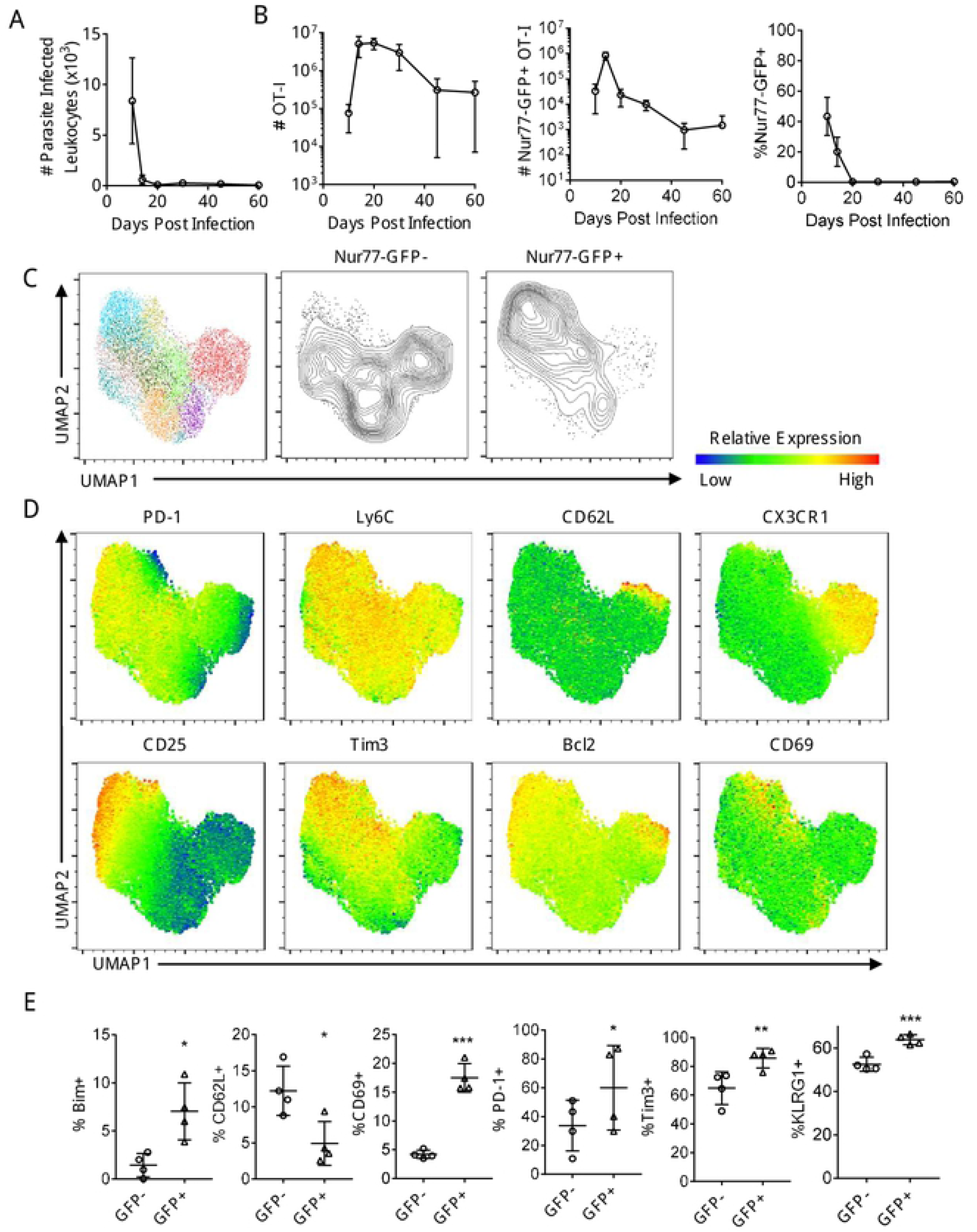
Infection induced kinetics and phenotype of splenic *T. gondii* specific CD8^+^ T cells. (A-B) OT-I were transferred i.v. into naive mice and infected with *T. gondii*-OVA. Mice were sacrificed between 10 and 60 dpi and spleens were analyzed (N=4-6 mice per time point). (C-E) High dimensional flow cytometry with UMAP and Phenograph clustering was performed on splenic OT-I at 10 dpi. P values based on Paired T-test; error bars indicate SEM.

To determine whether recent TCR stimulation is associated with a distinct phenotype, OT-I were analyzed by high-parameter flow cytometry using markers associated with effector and memory precursor populations (see Materials and Methods for full panel) at 10 dpi. Phenograph clustering indicated that these T cells that had not seen recent TCR appeared as five major clusters, one characterized by the expression of CX3CR1 and CD62L and another associated with reduced expression of activation markers such as CD25 (IL-2Rα) and PD-1 (Figure 2C). In contrast, a majority of GFP^+^ OT-I were defined by expression of CD25, PD-1, Bim, and Bcl-2, but were low for CX3CR1 and CD62L (Figure 2D, E). Additionally, GFP^+^ OT-I were more likely to be positive for Ki67 (a marker of entry into cell cycle) and KLRG1 (an inhibitory receptor expressed by effector T cells) (Figure 2E). Thus, at the peak of the T cell response in the periphery, recent TCR activation is most closely associated with those T cells that express markers of an effector population, but there was a population of effector memory precursor cells (CX3CR1^+^KLRG1^+^) that appeared less likely to receive TCR signals.

### Secondary TCR activation does not impact on ability to form memory

Previous studies have highlighted the impact of TCR activation on the formation of memory populations and the observation that the majority of GFP^-^ OT-I were KLRG1^+^ CX3CR1^+^, whereas the GFP^+^ OT-I were largely KLRG1^+^ CX3CR1^-^ (Figure 3A) suggested that these populations may have different memory potential. To determine if recent antigen stimulation affected the ability to generate memory cell populations, GFP^-^ and GFP^+^ OT-I were sorted from the spleen of day 10 *T. gondii*-OVA infected mice and transferred into naive mice (Figure 3B, C). This process did not lead to infection, based on the absence of tdTomato^+^ cells and lack of GFP expression in the transferred OT-I (Supplementary Figure 2A, B). Nevertheless, 14 days post transfer, OT-I were present in the spleens of these mice but there was no difference in the number of OT-I recovered from the two experimental groups (Figure 3D). In addition, despite differences in the expression of CX3CR1 prior to transfer, the memory cell populations in the mice were all characterized by high expression of CX3CR1 and KLRG1 (Figure 3E), as well as other effector and memory markers such as CXCR3 (Supplementary Figure 2C). These data suggest the population of TCR stimulated cells at day 10 retain the ability to give rise to an effector memory population characterized by co-expression of KLRG1 and CX3CR1.

**Fig 3.**
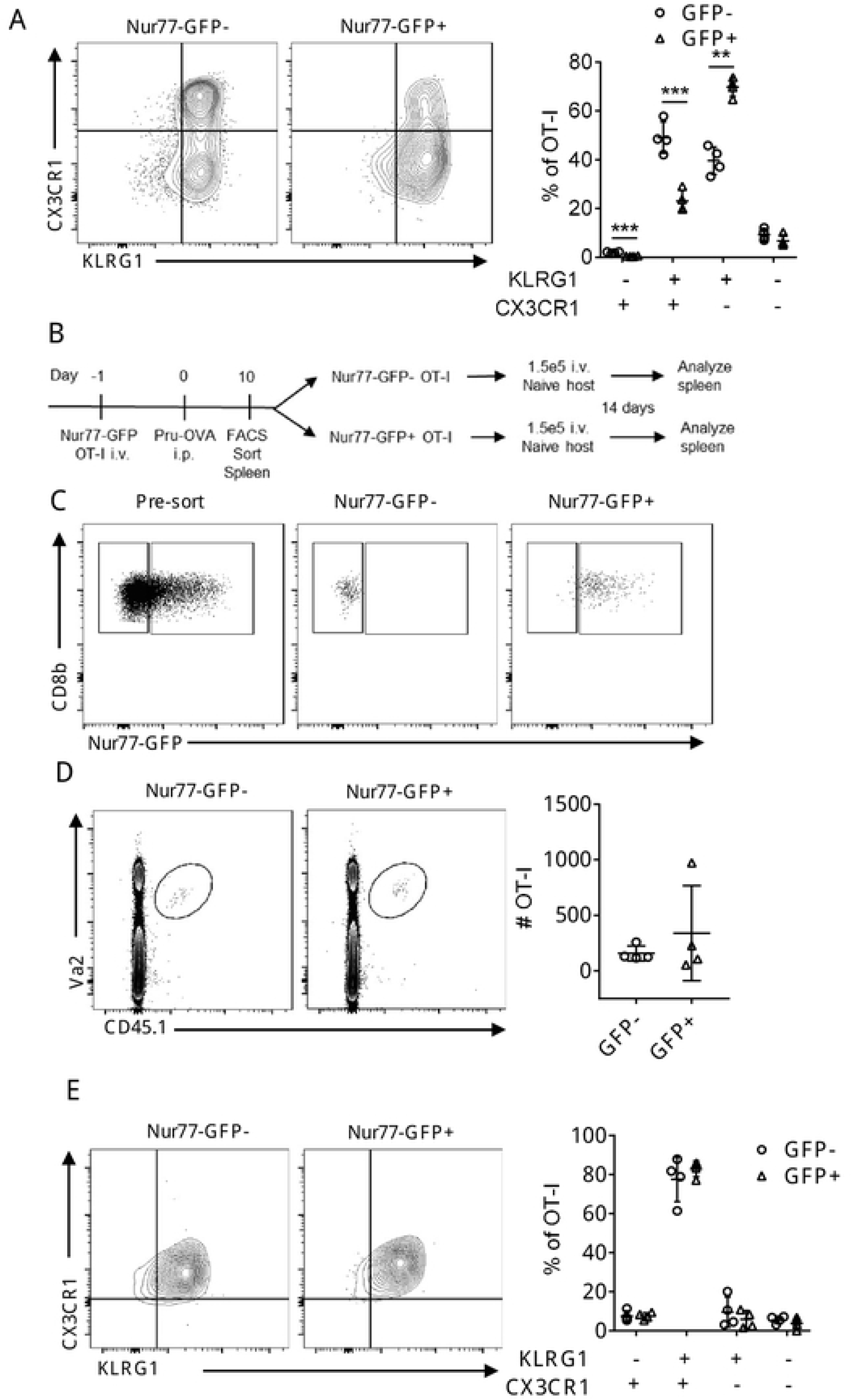
CD8^+^ T cell memory potential retained in Nur77-GFP^+^ OT-I. (A) OT-I from spleen of *T. gondii*-OVA 10 dpi. (B-C) OT-I from spleen of T. gondii-OVA 10 dpi were sorted by Nur77-GFP expression and transferred into naive mice. Mice were sacrificed 14 days post transfer (N=4 mice). (D) Identification of OT-I from mice receiving either Nur77-GFP^+^ or Nur77-GFP^-^ OT-I. (E-F) Characterization of OT-I from recipient mice. P values based on Students T-test with Bonferroni-Dunn correction for multiple comparisons; error bars indicate SEM.

### Influence of TCR stimulation on T cell responses in the CNS

In B6 mice, *T. gondii* enters the CNS as a tachyzoite and foci of parasite replication associated with areas of inflammation are readily apparent. As T cell responses lead to local parasite control, the ability to form tissue cysts (that contain bradyzoites) within neurons allows persistent infection, although periodic reactivation of cysts can lead to areas of tachyzoite replication. Indeed, parasites were detected in the CNS at 10 dpi and the number of tachyzoite infected leukocytes declined between days 20 and 30, although total parasite burden remained high through 40 dpi (Figure 4A). OT-I were recruited to the brain between 10 and 14 dpi (Figure 4B) when ∼40% of these cells expressed GFP which decreased to ∼10-20% of OT-I at 2 months post infection (Figure 4B). The high percentage of GFP^+^ cells in the CNS at 10 dpi is at a time point when T cells enter the CNS but infected endothelial cells in the vascular compartment are present [44]. To distinguish the OT-I T cells in the vascular compartment from those that had crossed into the parenchyma of the brain, fluorophore conjugated αCD45 was administered i.v. 3 minutes prior to sacrifice to label vascular resident cells and these populations were compared with those in whole blood. At 10 dpi in the whole blood ∼20% of OT-I were GFP^+^, with a rapid decline in GFP expression by 14 dpi (Figure 4C) which correlated with the clearance of parasites from circulation. In the CNS at 10 dpi, ∼30% of OT-I were vascular and ∼40% of these vascular OT-I expressed Nur77-GFP (Figure 4C). By 20 dpi, this frequency of brain vascular OT-I had dropped to <5% (Figure 4C) and expression of GFP in these OT-I cells was <20% (Figure4C). At 10 dpi when OT-I are entering the CNS, vascular associated Nur77-GFP+ OT-I were positive for CX3CR1 (Figure 4D), a chemokine receptor involved in T cell trafficking.

**Fig 4.**
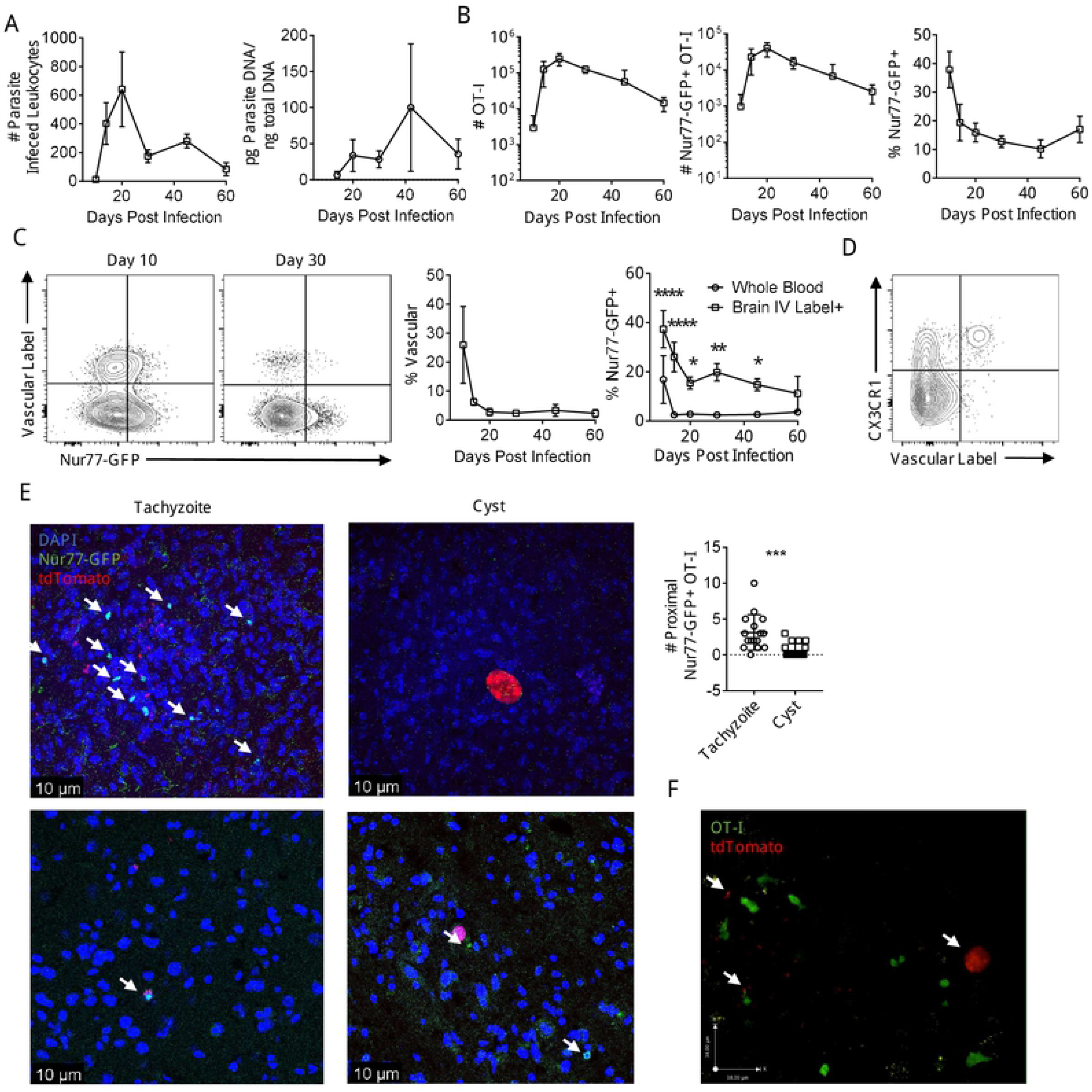
Infection induced kinetics of CNS *T. gondii* specific CD8^+^ T cells. (A-B) Kinetics of infection and OT-I in the brain following infection with *T. gondii*-OVA (N=4-6 mice per time point). (C) Mice were given i.v. dose of fluorophore conjugated αCD45 prior to sacrifice to label vascular cells. OT-I from whole blood were compared to OT-I from the vascular labeled fraction of the brain. P values based on Two-Way ANOVA; error bars indicate SEM. (D) Nur77- GFP+ OT-I from the CNS of *T. gondii*-OVA infected mice at 10 dpi. (E) Brain sections were stained for CD45.1 (white) to identify OT-I, anti-GFP (green) to identify recently TCR stimulated OT-I, DAPI (blue) to identify nuclei, and were infected with *T. gondii*-OVA expressing tdTomato (red). Number of Nur77-GFP+ OT-I surrounding parasites were counted. Arrows indicate Nur77-GFP^+^ OT-I. (F) *Ex vivo* imaging of constitutive GFP expressing OT-I (green) in brain explants of *T. gondii*-OVA (red) infected mice. Arrows indicate *T. gondii* infection. P values based on Students T-test; error bars indicate SEM.

To determine if there was any differential association in the brain of GFP^+^ OT-I with areas of parasite replication, IHC was used to visualize fluorescent parasites, CD45.1^+^ OT-I T cells, and staining for GFP at day 28. OT-I were widely dispersed throughout the CNS and the majority were Nur77-GFP^-^, but Nur77-GFP^+^ OT-I were readily observed in association with areas of tachyzoite replication (Figure 4E). In addition, it was noted that a small number of GFP^+^ cells were detected in the vicinity of the parasite cyst stage (Figure 4E). Similar results could be detected using live imaging of infected mice, but in which the OT-I did not express the Nur77-GFP reporter, but rather expressed GFP under the control of the CD4 promoter, as previously described [56]. In this setting, congregation of OT-I around areas of tachyzoite replication, with OT-I that are actively infected by parasite identifiable were readily identified (Supplemental Movie 1). It was also notable that in this OT-I T cells that were non-motile and had a rounded appearance (two behaviors associated with TCR engagement) could be visualized in proximity to a cyst. Together, these data sets highlight that as the infection transitions to the chronic stage in the CNS there are enhanced levels of TCR activation that occurs at the interface between the brain and the vascular system, and in the parenchyma TCR activity appears largely restricted to sites of parasite replication.

### Impact of TCR engagement on CD8^+^ T cells during Toxoplasmic Encephalitis

To assess the impact of recent TCR stimulation on the CD8^+^ T cell population in the CNS, GFP^-^ and GFP^+^ populations of OT-I T cells from the brains of mice infected for 14 days were analyzed by RNA sequencing. Principal component analysis revealed that GFP^-^ and GFP^+^ OT-I formed distinct clusters (Supplementary Figure 3A) and over 2,000 genes were differentially regulated between these populations (adjusted P-value <0.05, logFC >0.3, Figure 5A). In GFP^-^ OT-I, transcripts for multiple transcription factors associated with memory precursor formation and longevity were upregulated (Figure 5B). The transcriptional profile of GFP^+^ OT-I was characteristic of effector T cells and associated with an enrichment of the surface receptors *Cx3cr1* and *Il2ra*, and transcriptional factors (such as *Zeb2* and *Irf8*), as well as those involved in negative regulation of T cell responses (*Egr2* and *Egr3*, Figure 5B). Moreover, an unbiased GSEA analysis revealed the top enriched gene sets of the GFP^+^ OT-I were associated with immune cell activation and anti-pathogen responses (Supplementary Figure 3B). As expected, the GFP^+^ OT-I were also enriched for gene sets associated with effector functions such as cytotoxicity and IFN-γ production (Figure 5C). This analysis also revealed enrichment of gene sets involved in the negative regulation of TCR mediated signaling (Figure 5C), indicating that an inhibitory feedback loop for TCR signaling was already engaged in those cells.

**Fig 5.**
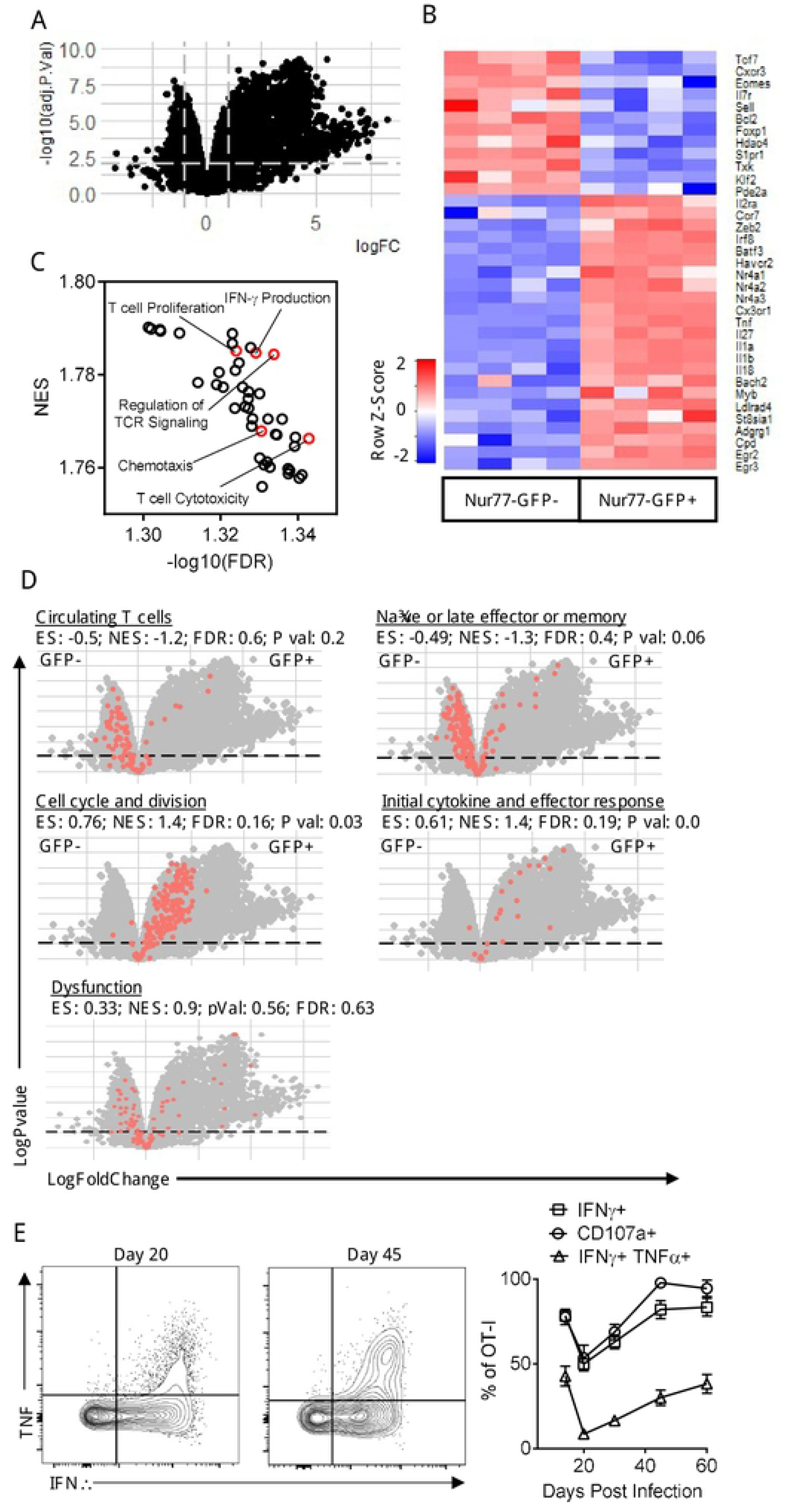
Transcriptional profiling of Nur77-GFP OT-I from CNS. (A-D) Nur77-GFP^-^ and Nur77-GFP^+^ OT-I from the brains of mice infected with *T. gondii*-OVA 14 dpi were transcriptionally profiled using RNA-sequencing. (A) Volcano plot of all transcripts. Lines indicate logFC > 0.03 and adjusted P-value > 0.05. (B) Heatmap of differentially expressed genes involved in CD8+ T cell activation, differentiation, memory formation, and effector function. (C) Top GSEA enriched pathways in Nur77-GFP^+^ OT-I compared to Nur77-GFP^-^ OT-I. (D) Gene sets involved in CD8+ T cell activation and differentiation (red dots). (E) OT-I from brains of *T. gondii*- OVA infected mice were restimulated with SIINFEKL peptide and cytokine and degranulation was assessed by flow cytometry.

To determine the differentiation status of GFP^-^ vs GFP^+^ OT-I, direct comparison of these subsets with gene signatures derived from LCMV specific CD8^+^ T cells at different points of infection [1] was performed. This analysis highlighted that GFP^-^ OT-I had greater abundance of transcripts connected with circulating T cells, as well as signatures associated with late effector CD8^+^ T cells (Figure 5D). Similar to the GSEA analysis (Figure 5C), the GFP^+^ OT-I were enriched for early effector molecules and anti-pathogen responses, as well as genes involved in cell division (Figure 5D). Although these T cells express multiple inhibitory receptors such as PD-1, there was no enrichment of a complete set of dysfunction related genes in either the GFP^-^ or GFP^+^ OT-I at this time point (Figure 5D). Previous studies have described the presence of a Trm-like population in the CNS during TE [33], and the Trm associated transcripts (*Cpd* and *Adgrg1)* were enriched in GFP^+^ OT-I, whereas transcripts that are decreased in Trm (*S1pr1* and *Klf2)* had lower abundance in GFP^+^ OT-I (Figure 5C). Furthermore, when the basal (i.e., not restimulated) ability of OT-I T cells in the brain to produce cytokines was assessed, GFP^+^ OT-I were 5 times more likely to be producing IFN-γ than GFP^-^ OT-I (Supplementary Figure 3C). For the OT-I T cells isolated from the brain between 14 and 60 dpi and stimulated with SIINFEKL peptide, the majority of these produced IFN-γ or TNF-α, and degranulated (based on CD107a expression, Figure 5E). These data indicate that TCR activation, rather than bystander activation, is the main stimulus required for effector function during TE and that at these time points there is no overt signature of T cell exhaustion.

Based in part on the transcriptional profiling, high parameter flow cytometry for a panel of markers of activation, memory, exhaustion, and transcription factors was employed to compare the Nur77-GFP OT-I T cell response in the spleen and brain at 2 and 6 weeks post infection. At 14 dpi OT-I phenotypes were distinct between the spleen and brain, with KLRG1 and CX3CR1 co-expression higher in the spleen than brain (Figure 6A). Unsupervised clustering of the brain OT-I at this time point identified 11 unique populations (Figure 6B). There were 7 clusters that were independent of recent TCR activation, and 4 clusters associated with Nur77-GFP expression (Figure 6B). Clusters that contained the GFP^+^ OT-I were also enriched for expression of KLRG1, CX3CR1, Tim3, CD25, Ki67 and Bcl-2 (Figure 6C, D). One of the most notable differences is illustrated by the GFP^+^ OT-I T cell co-expression of KLRG1 and CX3CR1 that are characteristic of effector T cells (Figure 6E).

**Fig 6.**
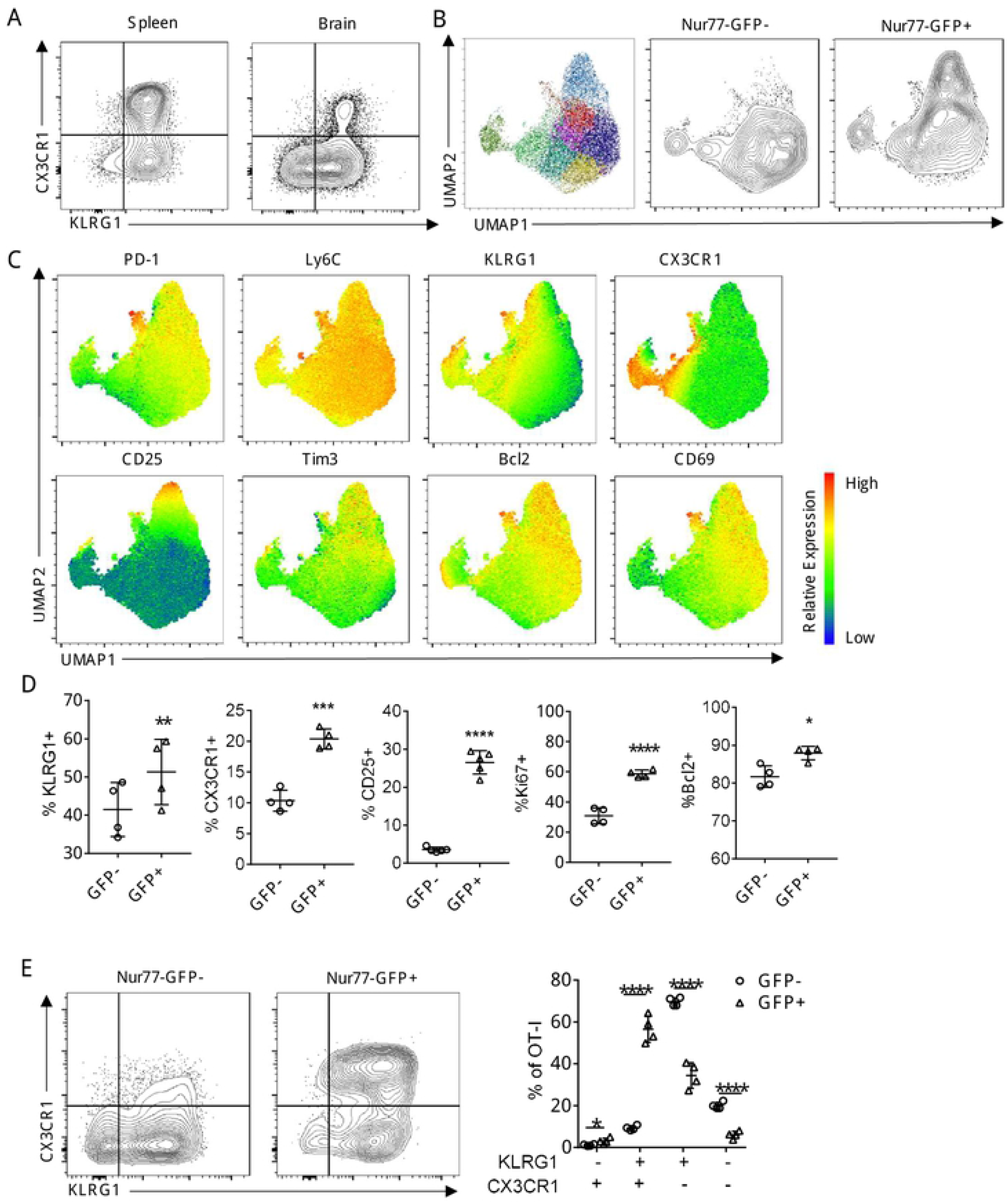
Infection induced phenotype of CNS *T. gondii* specific CD8^+^ T cells during early infection. (A) Phenotype of OT-I in spleen and brain of *T. gondii*-OVA infected mice 14 dpi. (B-E) High dimensional flow cytometry with UMAP and Phenograph clustering was performed on brain OT-I at 14 dpi. P values based on Paired T-test; error bars indicate SEM.

At 10 dpi, the splenic OT-I were a heterogenous population based on KLRG1 and CX3CR1 (Figure 3A) but by 6 weeks post infection, there was a more homogeneous (>98%) population of OT-I that maintained a KLRG1^+^CX3CR1^+^ phenotype (Figure 7A). In contrast, in the brain, the vast majority of OT-I were not positive for CX3CR1 and varied in KLRG1 expression (Figure 7A). In the CNS, segregation by UMAP of the OT-I from 2 and 6 weeks post infection were markedly different phenotypically (Figure 7B). At the 6 week timepoint, approximately 15% of OT-I in the brain were Nur77-GFP^+^ (Figure 4B). Brain OT-I formed 9 unique clusters (with Nur77-GFP excluded from the UMAP analysis), a subset of GFP^+^ OT-I clustered differently than GFP^-^ OT-I by UMAP (Figure 7C). As in the early phase of the T cell response in the brain, KLRG1, Bcl-2, CD25, and Ki67 expression were higher on GFP^+^ OT-I (Figure 7D, E) and recent TCR activation was still strongly predictive of IFN-γ expression (Figure 7F). These data demonstrate that TCR stimulation contributes to the heterogeneity seen in the CD8^+^ T cell population, and there is still considerable variation in T cell phenotypes unexplained by recent TCR stimulation.

**Fig 7.**
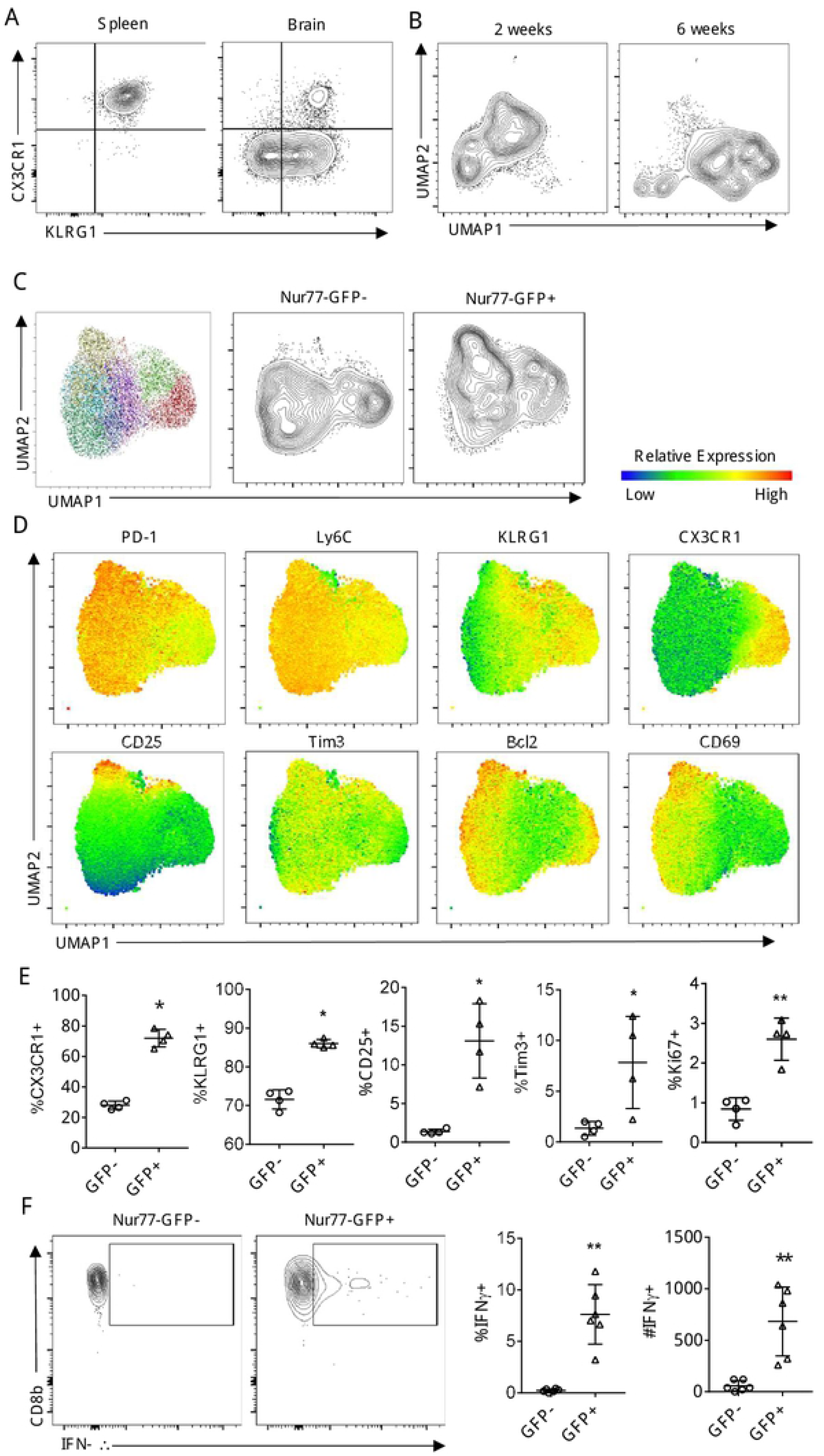
Infection induced phenotype and function of CNS *T. gondii* specific CD8^+^ T cells during chronic infection. (A) High dimensional flow cytometry with UMAP of OT-I from brains of mice infected with *T. gondii*- OVA at 14 dpi and 6 weeks post infection. (B) Phenotype of OT-I in spleen and brain of *T. gondii*- OVA infected mice 6 weeks post infection. (C-E) High dimensional flow cytometry with UMAP and Phenograph clustering was performed on brain OT-I at 6 weeks post infection. (F) Endogenous levels of IFN-γ expression in Nur77-GFP- and Nur77-GFP+ OT-I following 4-hour culture with protein transport inhibitor. P values based on Paired T-test; error bars indicate SEM.

### Impact of parasite replication on TCR on TRM CD8^+^ T cell phenotype

The transcriptional profiling at day 14 indicated the presence of a Trm signature, consistent with the idea that continued local antigen stimulation may promote or maintain the formation of local Trm cells [64]. Indeed, at 6 weeks post infection, more than 50% of the GFP^+^ OT-I coexpressed CD69 and CD103, suggesting that recent TCR activation promoted Trm (Figure 8A). To address how antigen levels impact OT-I heterogeneity, *T. gondii*-OVA infected mice were treated with sulfadiazine, a drug that inhibits tachyzoite replication, from days 21 to 49 post infection, and OT-I were subjected to high dimensional flow cytometry. As expected, based on the detection of tdTomato and PCR, treatment with sulfadiazine reduced parasite burden in the brain (Figure 8B), accompanied by a reduction in the percent of GFP^+^ OT-I, as well as the level of reporter expression per cell (Figure 8C). The OT-I isolated from the brains of sulfadiazine treated mice had partial differential clustering on UMAP (Figure 8D), and the most striking impact on this population was the loss of CX3CR1 expression by the GFP^+^ OT-I compared to vehicle treated mice (Figure 8E, F). Surprisingly, there was no significant change in the frequency or number of CD69^+^ CD103^+^ OT-I in the CNS (Figure 8E, G). These data sets establish that this population of Trm is not dependent on active parasite replication and suggests that antigen may influence Trm phenotype at an earlier stage of activation.

**Fig 8.**
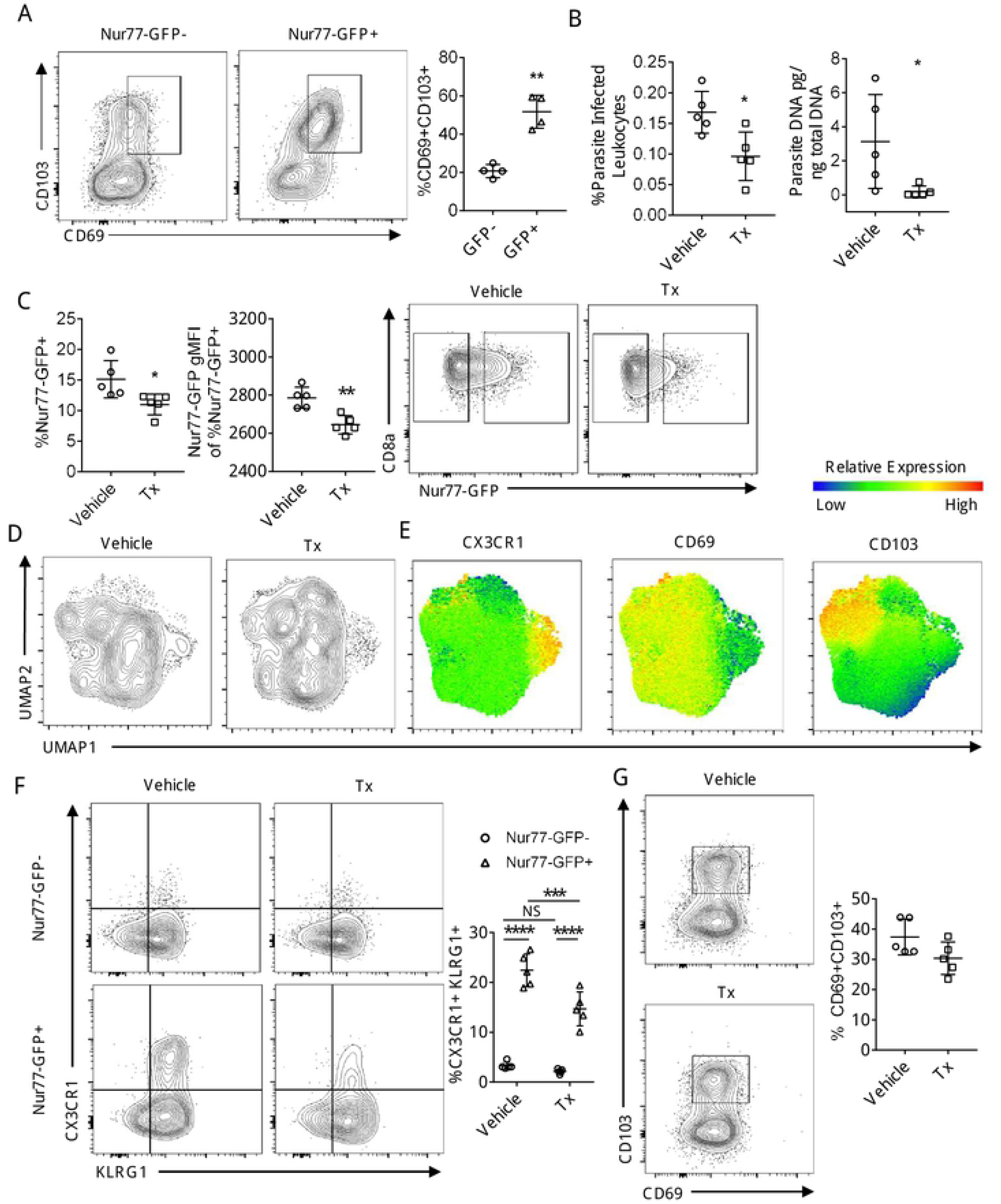
Impact of parasite replication of CNS CD8^+^ T cell phenotype. (A) Trm phenotype of OT-I at 6 weeks post infection in the brain. P values based on Paired T-test; error bars indicate SEM. (B-G) *T. gondii*-OVA infected mice were treated with sulfadiazine from 3 weeks to 7 weeks post infection and parasite burden and OT-I phenotype was assessed at 7 weeks. P values based on Students T-test (B-C) or Two-Way ANOVA (F); error bars indicate SEM.

## Discussion

Our previous studies have shown that during TE, parasite-specific CD8^+^ T cells are highly motile and based on their patterns of migration a mathematical model was developed for the ability of these cells to find infected targets [65]. In that report it was unclear what proportion or how frequently the parasite specific CD8^+^ T cells in the periphery or CNS re-encounter antigen and how this might influence effector function or fate. Here we show that TCR engagement in the periphery and the CNS correlates with levels of infection in these compartments and was as high as ∼40% but fell to 10-15% in the chronic stage of infection in the CNS. Given that this Nur77 reporter only provides a snapshot of how many T cells have seen antigen within a narrow time window it appears that the frequency with which CD8^+^ T cells encounter cognate antigen is high. Similarly, studies utilizing calcium flux reporters in the CNS of mice with high VSV titers concluded that up to 30% of pathogen specific CD8^+^ T cells were being stimulated by antigen over a short time period [24]. Thus, despite the CNS being a tissue canonically considered immune privileged there are robust levels of antigen presentation and T cell activation that is required for local pathogen control.

While initial TCR stimulation is necessary for T cell priming, few studies have distinguished the role of subsequent antigen exposure on the function and fate of activated effector and memory CD8^+^ T cells *in vivo*. Many cytokines and transcription factors contribute to the development of T cell heterogeneity, and during toxoplasmosis these include the cytokines IL-7 and IL-12 [66, 67], as well as transcription factors T-bet, c-Rel, BLIMP1, STAT1, and Bhlhe40 [68–73], yet it is unclear whether secondary antigen encounter and TCR activation alters CD8^+^ T cell fate decisions. The studies presented here show that TCR engagement of activated CD8^+^ T cells in a secondary site of infection correlated with profound transcriptional changes and expression of markers of effector T cells (for example co-expression of KLRG1 and CX3CR1 and cytokine production). It has been proposed that TCR signal strength and levels of inflammation during priming influence the balance between effector and memory CD8^+^ T cells [63, 74]. However, in the transfer studies performed here recent TCR signals did not appear to impact the ability to form a memory CD8^+^ T cell population. Likewise, the presence of Trm CD8^+^ T cells in the CNS have been described for other infections [75] and TCR signal strength and duration influences the establishment and function of these Trm populations [76, 77]. A Trm population has also been described during TE [33] and the studies presented here revealed that the OT-I Trm-like cells received frequent TCR stimulation. However, while the absence of active parasite replication resulted in reduced TCR engagement there was no significant change in the frequency or number of these Trm cells in the CNS. Thus, both approaches that addressed the impact of secondary TCR activity on memory populations suggest that recent antigen encounter does not alter these fates and fit with the concept that Trm progenitors are generated early in effector responses [75, 78].

The CNS represents a unique site of inflammation during infection, and the ability of *T. gondii* to establish a persistent infection in this tissue provides the opportunity to study how TCR signals intersect with different phases of the T cell response in the brain. For example, it has been suggested that following challenge with viral or bacterial infections that pathogen replication in the brain is not required for resident CD8^+^ T cell populations to populate the CNS [79]. However, during toxoplasmosis, endothelial cells of the blood brain barrier are infected [44] and the data that associates CX3CR1 (a chemokine receptor responsible for leukocyte entry into the CNS [80]) with TCR activation in the vascular compartment of the CNS implies that antigen stimulation at the blood brain barrier may induce T cell extravasation into the parenchyma. The use of the Nur77-GFP reporter as a surrogate for TCR activation should provide a sensitive system to assess the host factors that contribute to compartment (vascular, endothelial, and parenchymal) specific T cell activity in the CNS required for T cell mediated protective immunity.

In certain cancers and infection, one of the effects of repetitive antigen stimulation on CD8^+^ T cells is the induction of T cell exhaustion, denoted by an increase in inhibitory receptors and a decrease in cytokine secretion and cytolytic capacity [81–83]. During TE, parasite-specific T cells express high levels of inhibitory receptors such as PD-1 and TIGIT and it has been suggested that these T cell populations may be exhausted [82–85]. Indeed, there is a report that PD-L1 blockade leads to enhanced CD8^+^ T cell responses during TE [83], whereas in a similar setting the absence of TIGIT does not augment the T cell response [84]. The impact of inhibitory receptors during TE is complicated by evidence that PD-1 supports the generation of Trm in the CNS in viral infections [86–88]. Nonetheless, in our studies IFN-γ expression was closely correlated with expression of Nur77-GFP, indicating that TCR-independent bystander cytokine-mediated activation of CD8+ T cells [89–91] is not a major contributor to the production of IFN-γ. Moreover, the functional and transcriptional analysis did not reveal any early signs of exhaustion in Nur77-GFP^+^ OT-I. Indeed, the magnitude of differentially expressed genes associated with TCR activation in the CNS was similar in scope to the transcriptional changes observed during priming of OT-I T cells in other models [1]. Many of these upregulated genes were transcription factors and regulators (*Irf8, Batf3, Bach2, Egr2*) known to direct CD8^+^ T cell function, indicating that subsequent antigen encounter may restructure transcriptional programs. These results also revealed six clusters that were not segregated by Nur77-GFP expression and underscore the need for additional studies to distinguish how priming affects diversity versus the TCR-independent signals that affect the evolution of the T cell response during TE.

There are several neurotropic infections that have distinct developmental states associated with latency and for *T. gondii* this is exemplified by the ability to form the tissue cyst. It has been proposed that the presence of this stage in neurons, a cell type known for low MHC-I expression [92], would be sequestered by the immune response. The idea that neurons can act as a refuge has been challenged by evidence that MHC-I presentation of tachyzoite antigens by neurons contributes to parasite control [58]. Imaging of brains from mice chronically infected with *T. gondii* had higher numbers of Nur77-GFP^+^ OT-I surrounding tachyzoites but as the infection in the brain progresses the decline in levels of TCR engagement correlated with parasite control and the switch from tachyzoite to bradyzoite. However, there were examples where Nur77-GFP^+^ OT-I were in close proximity to intact cysts but whether this is due to presentation by infected neurons, other glial populations, or infiltrating DC or monocyte populations that interact with infected neurons [56] is currently unknown. Whether the cyst stage is under direct immune surveillance is also unclear but the ability of bradyzoites to actively block IFN-γ signaling [47] implies the need for the parasite to evade an active response against the cyst stage [43, 93]. Currently, there is a paucity of host and parasite specific tools to dissect the ability of host cells to process and present antigens from the bradyzoite life-stage. Future experiments utilizing bradyzoite-specific secretion of model antigens combined with the Nur77-GFP reporter will be utilized to address this knowledge gap.

## Materials and Methods

### Mice and infection

Nur77-GFP reporter mice, C57BL/6 mice, CD45.1 C57BL/6 mice, and OT-I mice were purchased from Jackson Laboratories. DPE-GFP transgenic mice were originally obtained from Ulrich H. von Andrian (CBR, Harvard, Boston MA) and were crossed to OT-I mice. All mice were housed in a specific-pathogen free environment at the University of Pennsylvania School of Veterinary Medicine in accordance with federal guidelines and with approval of the Institutional Animal Care and Use Committee. The generation of CPS-OVA parasites and PRU-OVA parasites were described previously [28, 59].

Parasites were cultured and maintained by serial passage on human foreskin fibroblast cells in the presence of parasite culture media (DMEM, 20% medium M199, 10% fetal bovine serum (FBS), 1% penicillin-streptomycin, 25 μg/ml gentamycin) which was supplemented with uracil for culturing CPS strain parasites (final concentration of 0.2 mM uracil). Tachyzoites of each strain were prepared for infection by serial 26g needle passage and filtered through a 5-μm-pore-size filter. 1 day prior to infection, 5,000 congenically distinct naive splenic Nur77-GFP OT-I were transferred i.v. into naive mice. Mice were infected i.p. with 10^4^ live parasites. In indicated experiments, mice were treated with 250mg/L of sulfadiazine in drinking water for 4 weeks beginning at 3 weeks post infection.

For adoptive transfers of OT-I from infected mice to naive mice, OT-I were double sorted on a BD FACS Aria. Each naive mice received an i.v. transfer of 150,000 Nur77-GFP^-^ or Nur77-GFP^+^ pooled from 2 infected mice. Recipient mice were treated with sulfadiazine two days prior and 1 week following transfer to ensure *T. gondii* infection was not transferred.

### Parasite quantification by PCR

Approximately 10mg of tissue was snap frozen and DNA was extracted with Qiagen DNeasy Blood and Tissue Kit according to manufacturer’s protocol. Real-time PCR was conducted with SYBR Green master mix according to manufacturer’s protocol, with 500ng of template DNA. *T. gondii* DNA was amplified with forward (5’-TCCCCTCTGCTGGCGAAAAGT-3’) and reverse (5’- AGCGTTCGTGGTCAACTATCGATTG-3’) primers to the B1 gene. PCR was performed on an Applied Biosystems ViiA7 with the following cycle conditions: Hold phase: 2min 50C, 10min 95C; PCR phase (occurs 50x): 15s at 95C, 1min @60C.

### Tissue processing

Spleens were grinded through a 70uM filter and red blood cells lysed with ACK lysis buffer. Brains were chopped into 1mm sections and incubated at 37C for 1.5 hours with 250ug/mL Collagenase/Dispase and 10ug/mL DNase, passed through a 70uM filter, then leukocytes were isolated through 30% and 60% Percoll density centrifugation.

### Quantification of T cell cytokine production

1x10^6^ brain mononuclear cells were cultured *ex vivo* at 37C for 4 hours in RPMI supplemented with 10% FBS. For restimulations, 1uM of peptide was added with Protein Transport Inhibitor Cocktail. For quantification of endogenous levels of cytokine production, only Protein Transport Inhibitor Cocktail was added.

### Bone marrow macrophage culture

Bone marrow macrophages were isolated and cultured for 7 days with 30% L929 media in DMEM with 10% FBS, with media replenishing on day 4 after isolation. For peptide stimulations, 1uM of peptide was added to macrophages for 20 minutes and excess peptide was removed by washing 2 times. For parasite infection, a parasite to macrophage infection ratio of 2:1 was used, and excess parasites were removed by washing 2 times. CD8^+^ T cell were isolated by MACS bead enrichment. Macrophages and T cells were cultured at a 1:2 ratio.

### Flow cytometry

Staining of extracellular antigens was performed following Fc receptor blocking with Rat IgG and anti-mouse CD16/CD32. Intracellular antigens were stained following fixation and permeabilization using the Foxp3 Transcription Factor Staining Buffer Set. All samples were run on a BD FACSymphony A5 or BD FACSCanto and analyzed on FlowJo v10 software (FlowJo, Ashland, OR). UMAP and Phenograph were utilized in FlowJo for dimensionality reduction and unsupervised clustering.

### Imaging

Brains were fixed in 4% paraformaldehyde overnight, followed by 15% and 30% sucrose in PBS overnight. 12uM sections were stained with AF488 anti-GFP and AF647 anti-CD45.1 in 0.1% Triton-X100 and 0.05% Tween-20 in PBS. Slides were sealed with Prolong Diamond Antifade Mountant with DAPI. Slides were imaged at 60x magnification on a Leica SP5-II (Lecia, Wetzlar, Germany).

Live imaging of brain explants was performed as previously described [65]. Briefly, mice were infected with PRU-OVA parasites expressing tdTomato. Four weeks following infection, mice received 2x10^6^ activated DPE-GFP-OT-I i.v., and brains were imaged one week following transfer of DPE-GFP-OT-I. Imaging was performed on an Lecia SP5 multiphoton.

### RNA sequencing

Nur77-GFP- and Nur77-GFP+ OT-I were double sorted as described above. Cells were collected in RPMI with 20% FBS and resuspended in Buffer RLT Plus. mRNA was isolated using RNeasy Micro Plus Kit and RNA quality was evaluated by High Sensitivity RNA Screen Tape on an Agilent 4200 TapeStation. cDNA library was prepared with Takara SMART-Seq HT kit and adapters ligated with Nextera XT DNA Library Preparation Kit and Index Kit v2 according to manufacturers’ protocols. AMPure XP beads were used for primer cleanup. 75-base pair reads were sequenced on a NextSeq 500 machine (Illumina, San Diego, CA)) according to manufacturer protocol. Kallisto was utilized to pseudo-align genes to version 75 of the mouse reference genome GRCm38. Transcripts that were represented in fewer than 3 of 4 samples in a group were excluded from analysis. The limma package in R was used to generate normalized counts per million, and differential genes (adjusted P-value>0.05) were identified with a log fold change cutoff of 0.3. The heatmaply package was used for heatmap generation. Gene set enrichment analysis was performed with GSEA software (v4.0.3, Broad Institute) with gene sets containing less than 10 genes excluded. Enrichment plots were generated in Cytoscape (v3.7.1) with an adjusted P-value cutoff of 0.05 and false discovery rate cutoff of 0.1 (pipeline adapted from [94]). Data is deposited in GEO (accession# GSE188495).

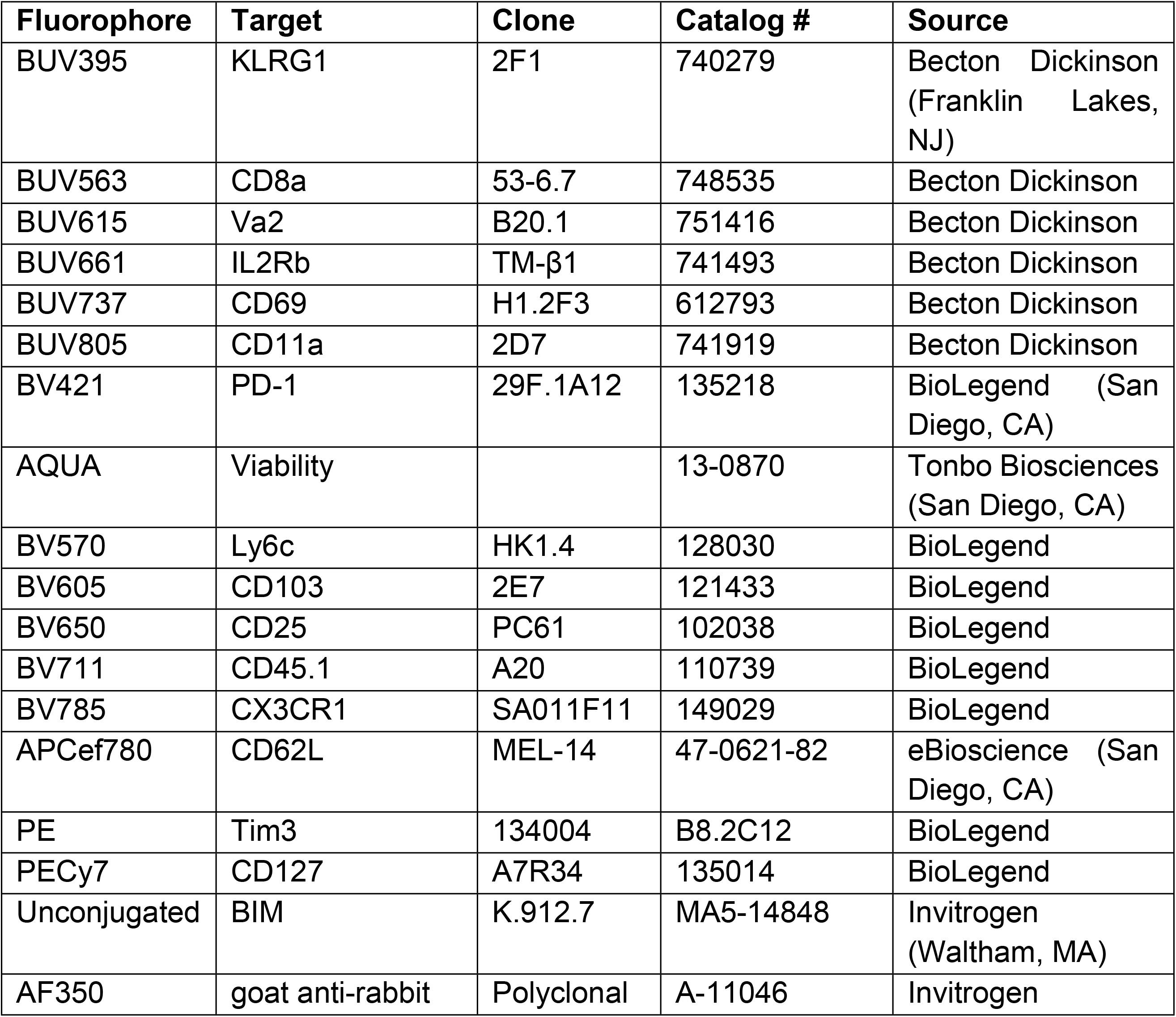
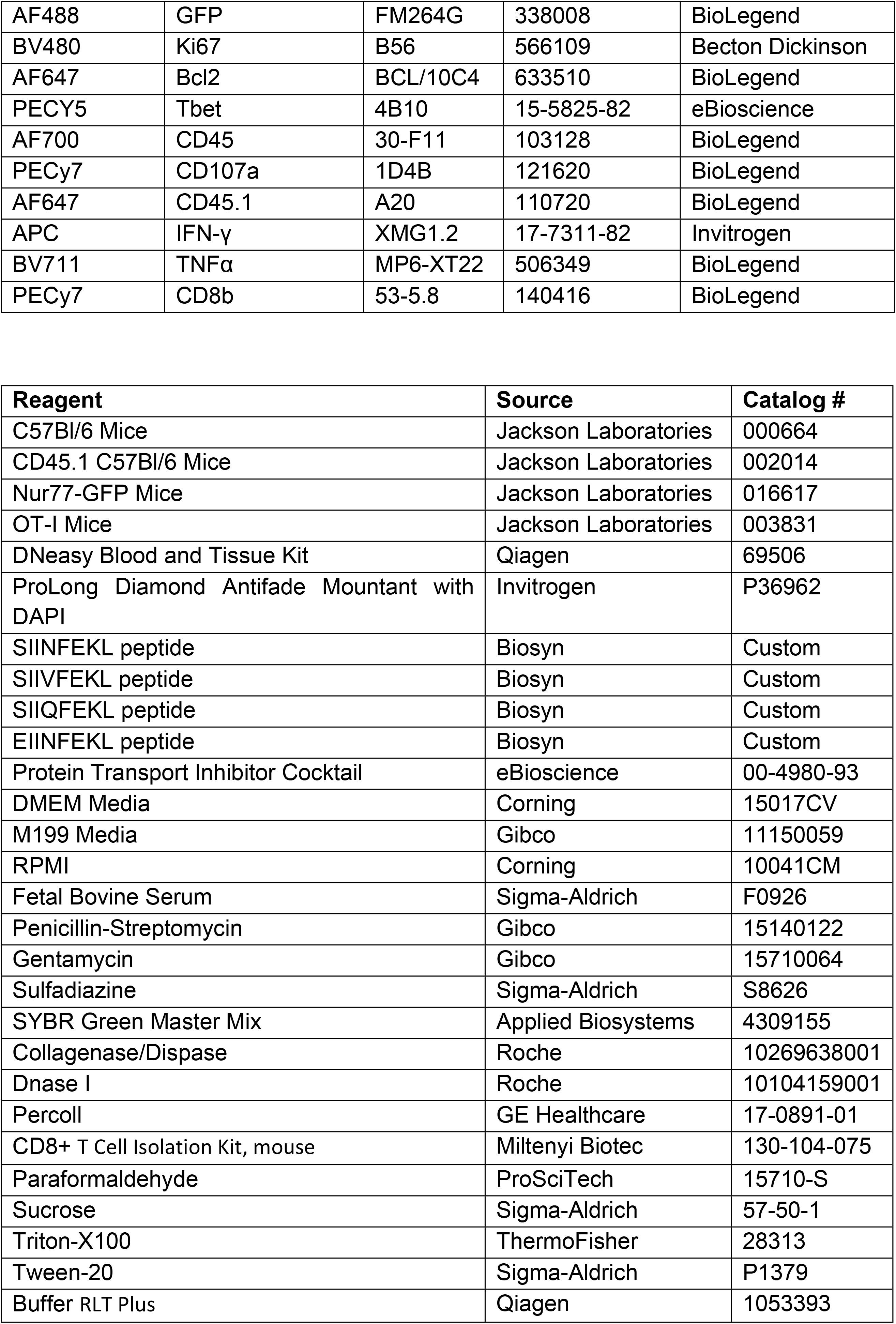
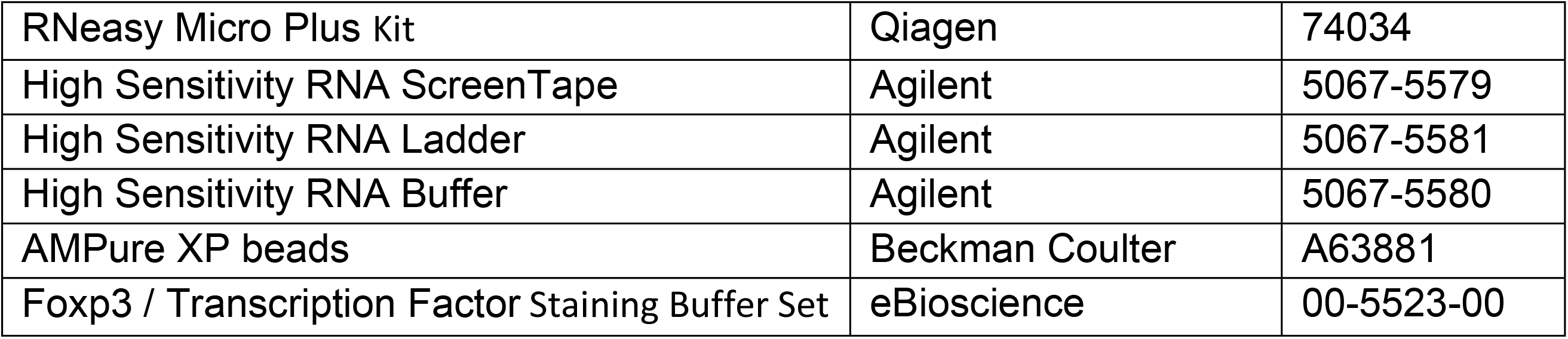

| Fluorophore | Target | Clone | Catalog # | Source |
| --- | --- | --- | --- | --- |
| BUV395 | KLRG1 | 2F1 | 740279 | Becton Dickinson (Franklin Lakes, NJ) |
| BUV563 | CD8a | 53-6.7 | 748535 | Becton Dickinson |
| BUV615 | Va2 | B20.1 | 751416 | Becton Dickinson |
| BUV661 | IL2Rb | TM-β1 | 741493 | Becton Dickinson |
| BUV737 | CD69 | H1.2F3 | 612793 | Becton Dickinson |
| BUV805 | CD11a | 2D7 | 741919 | Becton Dickinson |
| BV421 | PD-1 | 29F.1A12 | 135218 | BioLegend (San Diego, CA) |
| AQUA | Viability |  | 13-0870 | Tonbo Biosciences (San Diego, CA) |
| BV570 | Ly6c | HK1.4 | 128030 | BioLegend |
| BV605 | CD103 | 2E7 | 121433 | BioLegend |
| BV650 | CD25 | PC61 | 102038 | BioLegend |
| BV711 | CD45.1 | A20 | 110739 | BioLegend |
| BV785 | CX3CR1 | SA011F11 | 149029 | BioLegend |
| APCef780 | CD62L | MEL-14 | 47-0621-82 | eBioscience (San Diego, CA) |
| PE | Tim3 | 134004 | B8.2C12 | BioLegend |
| PECy7 | CD127 | A7R34 | 135014 | BioLegend |
| Unconjugated | BIM | K.912.7 | MA5-14848 | Invitrogen (Waltham, MA) |
| AF350 | goat anti-rabbit | Polyclonal | A-11046 | Invitrogen |
| AF488 | GFP | FM264G | 338008 | BioLegend |
| BV480 | Ki67 | B56 | 566109 | Becton Dickinson |
| AF647 | Bcl2 | BCL/10C4 | 633510 | BioLegend |
| PECy5 | Tbet | 4B10 | 15-5825-82 | eBioscience |
| AF700 | CD45 | 30-F11 | 103128 | BioLegend |
| PECy7 | CD107a | 1D4B | 121620 | BioLegend |
| AF647 | CD45.1 | A20 | 110720 | BioLegend |
| APC | IFN-γ | XMG1.2 | 17-7311-82 | Invitrogen |
| BV711 | TNFα | MP6-XT22 | 506349 | BioLegend |
| PECy7 | CD8b | 53-5.8 | 140416 | BioLegend |

| Reagent | Source | Catalog # |
| --- | --- | --- |
| C57Bl/6 Mice | Jackson Laboratories | 000664 |
| CD45.1 C57Bl/6 Mice | Jackson Laboratories | 002014 |
| Nur77-GFP Mice | Jackson Laboratories | 016617 |
| OT-I Mice | Jackson Laboratories | 003831 |
| DNeasy Blood and Tissue Kit | Qiagen | 69506 |
| ProLong Diamond Antifade Mountant with DAPI | Invitrogen | P36962 |
| SIINFEKL peptide | Biosyn | Custom |
| SIIVFEKL peptide | Biosyn | Custom |
| SIIQFEKL peptide | Biosyn | Custom |
| EIINFEKL peptide | Biosyn | Custom |
| Protein Transport Inhibitor Cocktail | eBioscience | 00-4980-93 |
| DMEM Media | Corning | 15017CV |
| M199 Media | Gibco | 11150059 |
| RPMI | Corning | 10041CM |
| Fetal Bovine Serum | Sigma-Aldrich | F0926 |
| Penicillin-Streptomycin | Gibco | 15140122 |
| Gentamycin | Gibco | 15710064 |
| Sulfadiazine | Sigma-Aldrich | S8626 |
| SYBR Green Master Mix | Applied Biosystems | 4309155 |
| Collagenase/Dispase | Roche | 10269638001 |
| Dnase I | Roche | 10104159001 |
| Percoll | GE Healthcare | 17-0891-01 |
| CD8+ T Cell Isolation Kit, mouse | Miltenyi Biotec | 130-104-075 |
| Paraformaldehyde | ProSciTech | 15710-S |
| Sucrose | Sigma-Aldrich | 57-50-1 |
| Triton-X100 | ThermoFisher | 28313 |
| Tween-20 | Sigma-Aldrich | P1379 |
| Buffer RLT Plus | Qiagen | 1053393 |
| RNeasy Micro Plus Kit | Qiagen | 74034 |
| High Sensitivity RNA ScreenTape | Agilent | 5067-5579 |
| High Sensitivity RNA Ladder | Agilent | 5067-5581 |
| High Sensitivity RNA Buffer | Agilent | 5067-5580 |
| AMPure XP beads | Beckman Coulter | A63881 |
| Foxp3 / Transcription Factor Staining Buffer Set | eBioscience | 00-5523-00 |

**Supplementary Fig 1.**
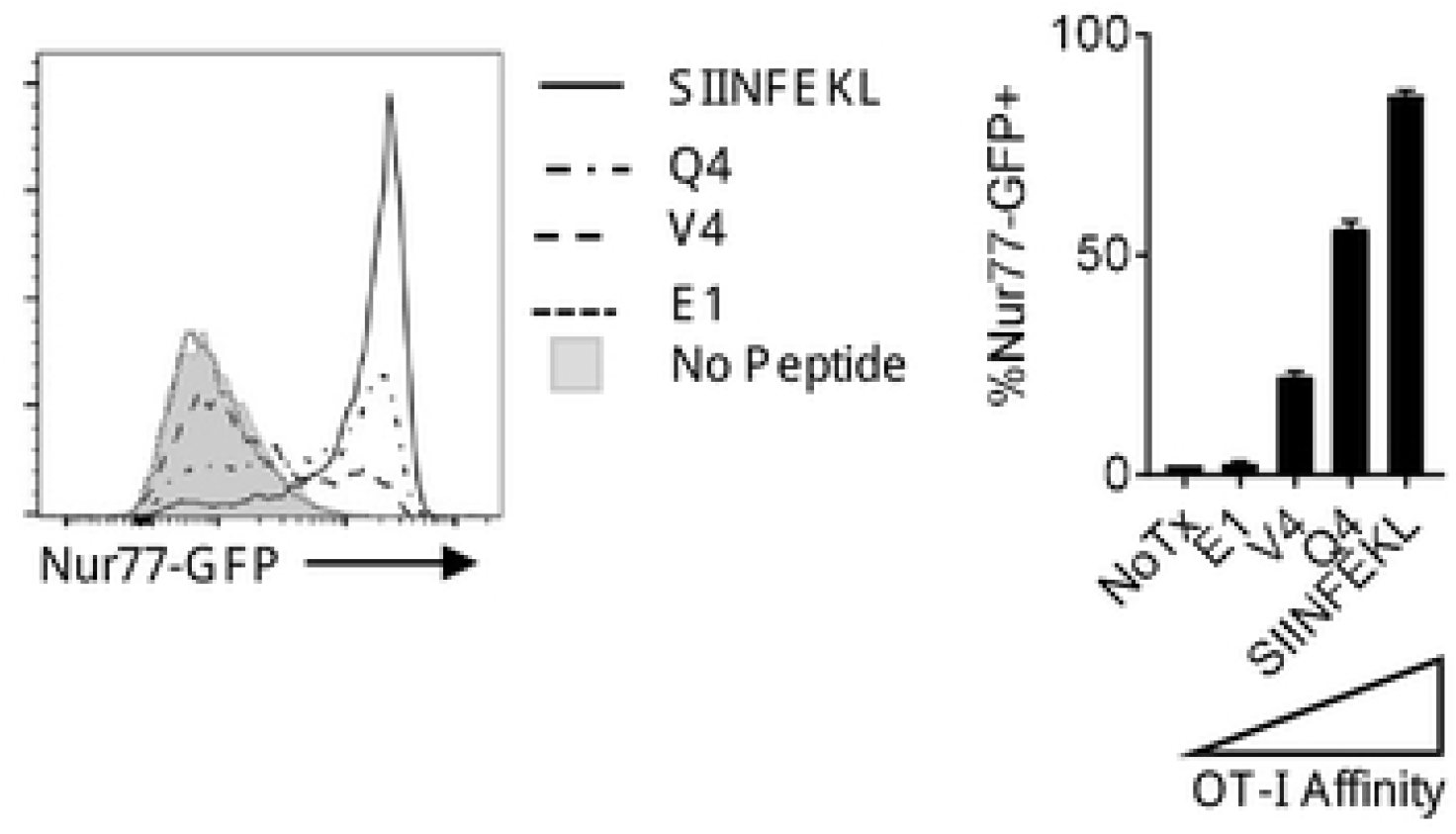
Effect of TCR signal strength on Nur77-GFP OT-I reporter activity. (A) Peptide pulsed splenocytes were incubated with Nur77-GFP OT-I and reporter activity was assessed at 3 hours.

**Supplementary Fig 2.**
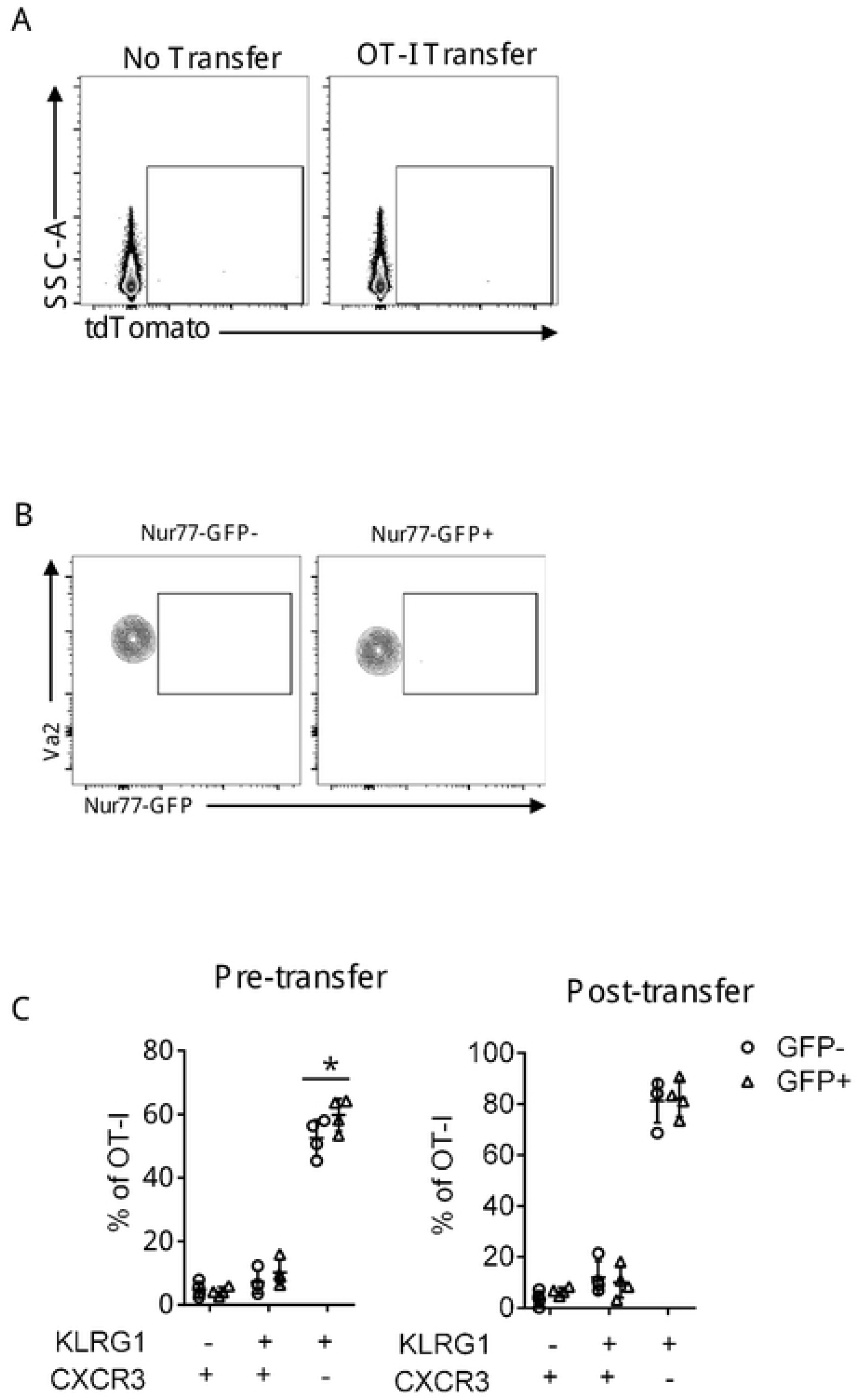
Transfer of Nur77-GFP OT-I from *T. gondii* infected mice into naive mice. Lack of infection at 14 days post transfer measured by (A) tdTomato expression of total splenocytes or (B) Nur77-GFP expression in transferred OT-I. (C) KLRG1 and CXCR3 expression of Nur77-GFP- and Nur77-GFP+ OT-I prior to transfer at 10 dpi and 14 days post transfer. P values based on Two-Way ANOVA; error bars indicate SEM.

**Supplementary Fig 3.**
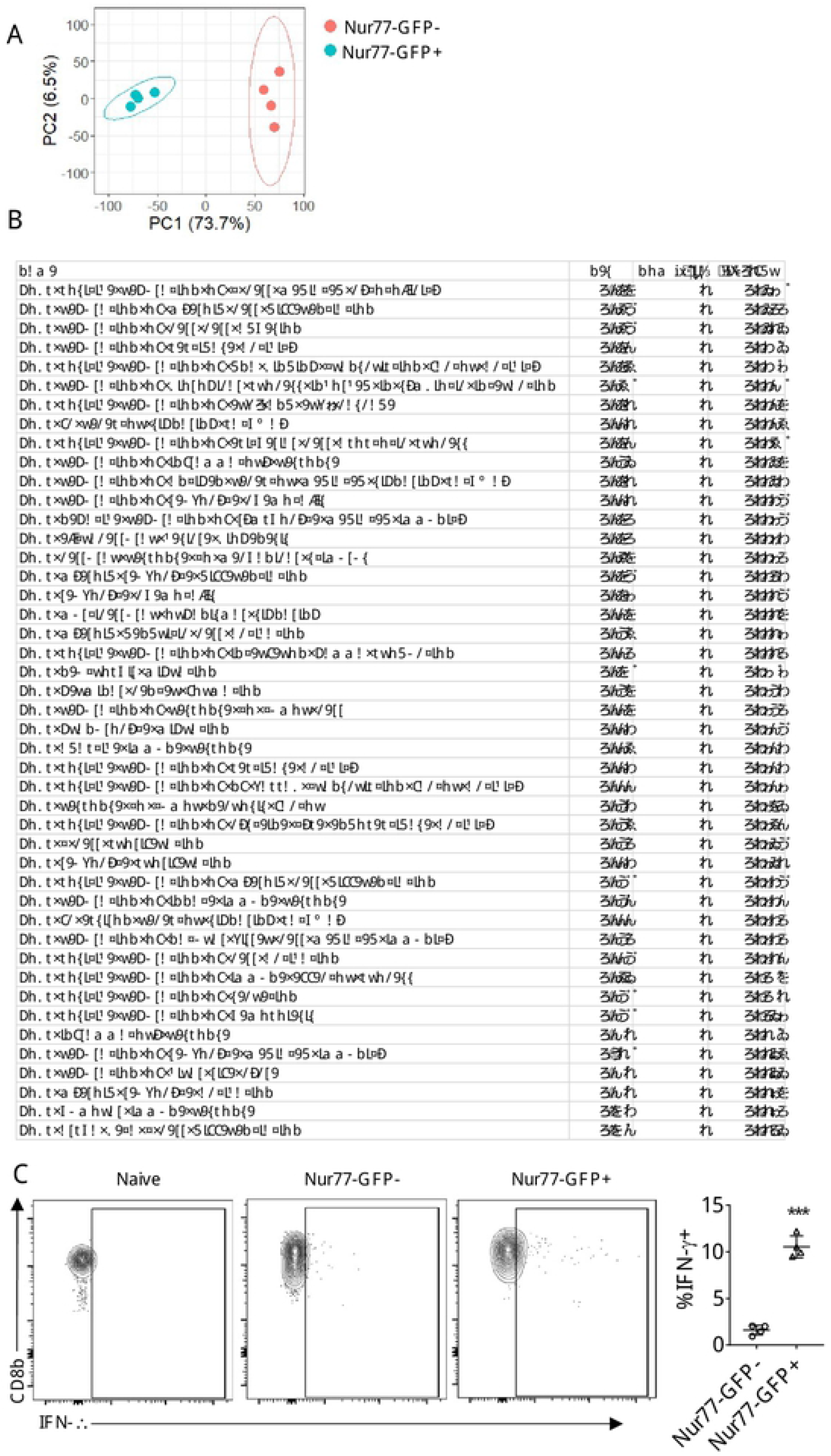
Transcriptional profiling of Nur77-GFP OT-I from CNS. (A) PCA plot of Nur77-GFP^-^ and Nur77-GFP^+^ OT-I. (B) Top GSEA gene sets with FDR>0.05 enriched in Nur77-GFP^+^ OT-I compared to Nur77-GFP^-^ OT-I. (C) Endogenous cytokine production of Nur77-GFP^-^ and Nur77-GFP^+^ OT-I from the CNS of *T. gondii*-OVA infected mice at 14 dpi.

Supplemental Movie 1. *Ex vivo* imaging of OT-I in the CNS of *T. gondii* infected mice.

Behavior of constitutive GFP OT-I T cells (green) in the brains of mice infected with *T. gondii*-OVA (red). OT-I were transferred into chronically infected mice and imaged one week post transfer. Live imaging of brain explants.

## Notes

### Competing Interest Statement

The authors have declared no competing interest.

